# Unadjuvanted intranasal spike vaccine booster elicits robust protective mucosal immunity against sarbecoviruses

**DOI:** 10.1101/2022.01.24.477597

**Authors:** Tianyang Mao, Benjamin Israelow, Alexandra Suberi, Liqun Zhou, Melanie Reschke, Mario A Peña-Hernández, Huiping Dong, Robert J. Homer, W. Mark Saltzman, Akiko Iwasaki

## Abstract

As the SARS-CoV-2 pandemic enters its third year, vaccines that not only prevent disease, but also prevent transmission are needed to help reduce global disease burden. Currently approved parenteral vaccines induce robust systemic immunity, but poor immunity at the respiratory mucosa. Here we describe the development of a novel vaccine strategy, Prime and Spike, based on unadjuvanted intranasal spike boosting that leverages existing immunity generated by primary vaccination to elicit mucosal immune memory within the respiratory tract. We show that Prime and Spike induces robust T resident memory cells, B resident memory cells and IgA at the respiratory mucosa, boosts systemic immunity, and completely protects mice with partial immunity from lethal SARS-CoV-2 infection. Using divergent spike proteins, Prime and Spike enables induction of cross-reactive immunity against sarbecoviruses without invoking original antigenic sin.

**One-sentence summary:** Broad sarbecovirus protective mucosal immunity is generated by unadjuvanted intranasal spike boost in preclinical model.

## Introduction

During the past two years of SARS-CoV-2 pandemic, we have seen the unprecedented development of highly effective vaccines, utilizing novel technologies including modified mRNA encapsulated in lipid nanoparticles (LNP) and replication deficient adenoviral vectors. Phase 3 clinical trials and subsequent post marketing vaccine effectiveness studies initially showed >90% vaccine efficacy against symptomatic disease (*1-3*). Additionally, early transmission studies showed decreased rates of transmission in household members of vaccinated individuals (*4, 5*). Unfortunately, more recent studies have demonstrated decreasing vaccine effectiveness in terms of asymptomatic infection as well as symptomatic and severe infections starting around 4 months post second dose with mRNA-LNP based regimens and earlier with other vaccines (*6, 7*). Furthermore, continued viral evolution with increasing immune evasiveness notably with Beta (B.1.351), Delta (B.1.617.2), and now Omicron (B.1.529) variants of concern (VOC), has also contributed to decreased vaccine effectiveness against COVID-19 (*8-10*). Not only have current vaccines become less effective at preventing SARS-CoV-2 infection, but they have also become less able to prevent viral transmission, which is likely due to multiple factors including increased viral transmissibility, viral immune evasion, waning vaccine effectiveness, and poor induction of respiratory mucosal immunity (*11*).

Currently approved SARS-CoV-2 mRNA-LNP-based and vector-based vaccines rely on intramuscular administration, which induces high levels of circulating antibodies, memory B cells and circulating effector CD4^+^ and CD8^+^ T cells in animal models and humans (*12-14*). However, parenteral vaccines do not induce high levels of potent antiviral immune memory at sites of infection such as tissue resident memory T cells (T_RM_) and B cells (B_RM_) as well as mucosal IgG and dimeric IgA (*15-17*). This is in contrast to SARS-CoV-2 infection in humans and mice where CD8^+^ T_RM_ are robustly induced (*15, 18*). Vaccines targeting the respiratory mucosa could address the shortcomings of parenteral vaccination, as recent preclinical assessments of intranasally delivered SARS-CoV-2 spike encoding adenoviral vectors have shown impressive mucosal immunogenicity as well as protection and reduced viral shedding in mice, hamsters, and nonhuman primates (*19-23*). Preclinical mucosal influenza vaccine studies have also shown that mucosal immunity can enhance protection against heterosubtypic challenge via CD8^+^ T_RM_ or dimeric IgA and may improve durability of immunity (*24, 25*).

While primary respiratory administration of vaccines induces potent mucosal immune responses, some studies have shown that priming systemically followed by intranasal boosting results in similar systemic immunity to systemic prime and boost regimens, but with the added benefit of enhanced mucosal immunity (*26, 27*). Most recombinant subunit vaccines administered via systemic priming followed by intranasal boosting or intranasal prime and boost require adjuvants to enhance immunogenicity. However, administration of vaccines to the respiratory tract in humans has proven difficult as vaccine associated adverse events within the respiratory tract are typically less tolerated than with systemic administration. Additionally, there have been cases of intranasal adjuvanted inactivated vaccine for seasonal influenza leading to non-respiratory adverse events (Bell’s palsy) in patients, which was hypothesized to be caused by adjuvant-mediated inflammation (*28*).

In the setting of waning immunity from parenteral vaccination regimens we assessed the immunogenicity and protection afforded by intranasal boosting (IN) SARS-CoV-2 spike. Here we describe a vaccination strategy that utilizes the strong systemic priming of mRNA-LNP based vaccine followed by IN boosting with either unadjuvanted spike protein or an immunosilent polyplex encapsulating mRNA in a preclinical model of COVID-19.

## Results

### IN boosting with unadjuvanted SARS-CoV-2 spike induces mucosal humoral immunity

To assess the potential of IN unadjuvanted subunit vaccine boosting for the development of respiratory tract mucosal immunity, we decided to harness the strong systemic immunogenicity of mRNA-LNP. We additionally benefited from extensive SARS-CoV-2 spike engineering which helps stabilize the protein in its prefusion confirmation with the addition of a C-terminal T4 fibritin trimerization motif, six proline substitutions (F817P, A892P, A899P, A942P, K986P, V987P), and alanine substitutions (R683A and R685A) in the furin cleavage site (*29*). These sets of mutations have been shown to significantly enhance immunogenicity and increase protein stability, some of which are used in current vaccines.

We vaccinated K18-hACE2 mice with 1 μg of mRNA-LNP (Comirnaty) by IM injection (Prime), followed 14 days later by 1 μg of recombinant unadjuvanted spike protein by IN administration (Prime and Spike). Additional control groups include K18-hACE2 mice that received IM Prime only and mice that received IN spike only at boosting. Mice were euthanized at day 21 or 28 (7- or 14-days post boosting) and assessed for the development of mucosal humoral immunity (Fig 1A).

**Fig 1.**
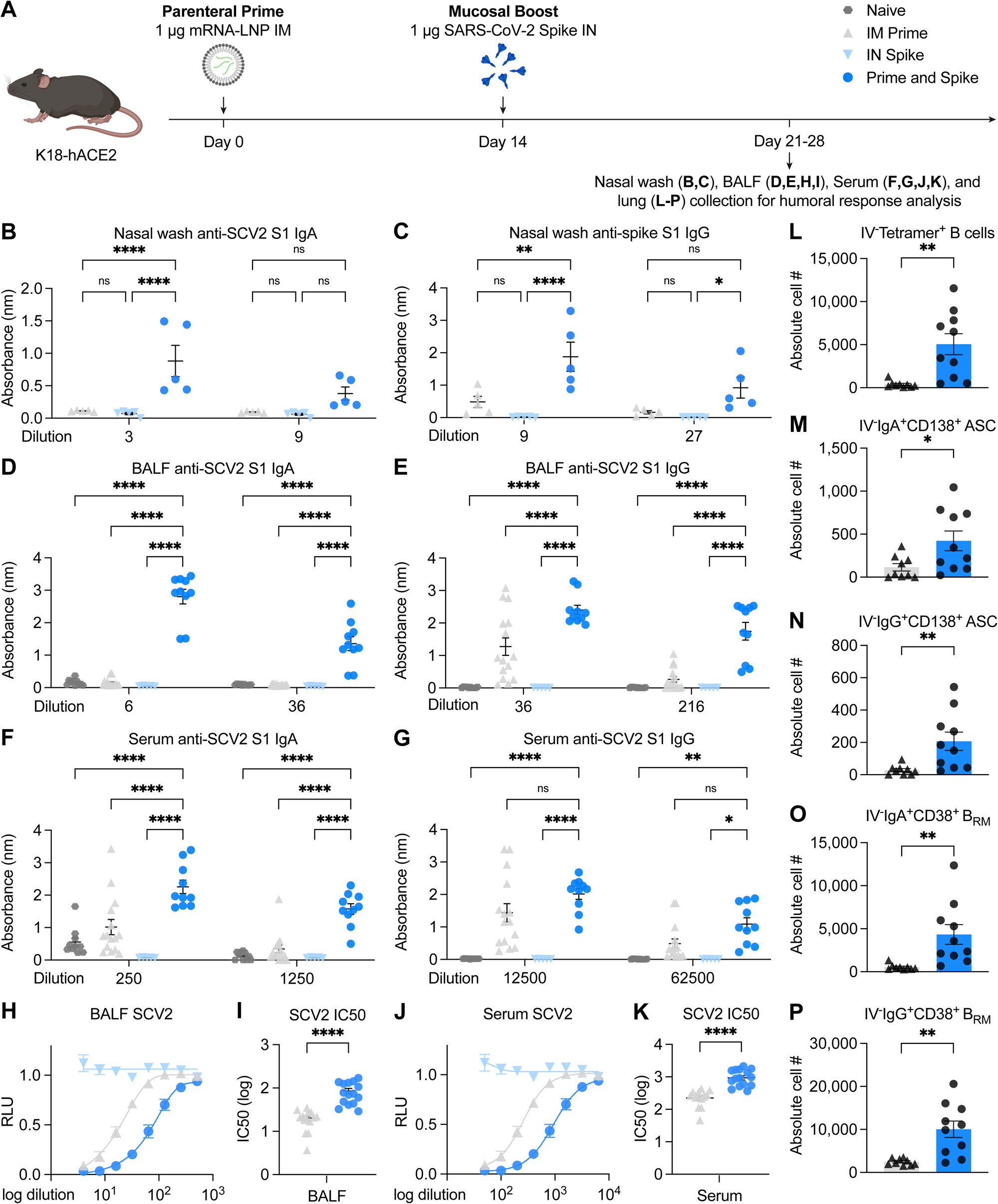
IN boosting with stabilized SARS-CoV-2 spike induces mucosal humoral memory. (**A**) Experimental schema: K18-hACE2 mice were intramuscularly (IM) immunized with 1 μg of mRNA-lipid nanoparticles (LNP) encoding full-length SARS-CoV-2 (SCV2) spike protein, followed by intranasal (IN) immunization with 1 μg of prefusion-stabilized, trimeric, recombinant SARS-CoV-2 (SCV2) spike protein 14 days following mRNA-LNP immunization. Fourteen (14) days post IN boost, serum, bronchoalveolar lavage fluids (BALF), and nasal washes were collected to assess binding and neutralizing antibody responses. Lung tissues were collected for extravascular B cell analysis. (**B-G**) Measurement of SCV2 spike S1 subunit-specific nasal wash IgA (**B**), nasal wash IgG (**C**), BALF IgA (**D**), BALF IgG (**E**), serum IgA (**F**), and serum IgG (**B**) in naïve mice, mice immunized with mRNA-LNP IM (IM Prime), mice immunized with the spike protein IN (IN Spike), or mice IM primed and IN boosted with spike (Prime and Spike). (**H-K**) Measurement of neutralization titer against SCV2 spike-pseudotyped vesicular stomatitis virus (VSV) in BALF (**H**,**I**) and serum (**J**,**K**). (**L-P**) Measurement of various extravascular (intravenous labeling antibody negative) B cell subsets, including RBD tetramer-binding B cells, IgA^+^ resident memory B cells (B_RM_), IgG^+^ B_RM_, IgA^+^ antibody secreting cells (ASC), and IgG^+^ ASC in lung tissues from IM Prime or Prime and Spike mice. Mean ± s.e.m.; Statistical significance was calculated by two-way ANOVA followed by Tukey correction (**B-G**) or student’s t-test (**I**,**K**,**L-P**); *P ≤ 0.05, **P ≤ 0.01, ***P ≤ 0.001, ****P ≤ 0.0001. Data are pooled from three independent experiments.

First, we assessed anti-SARS-CoV-2 spike S1 IgG and IgA in nasal wash, bronchoalveolar lavage fluid (BALF), and serum. We found that only mice that received Prime and Spike developed high levels of anti-SARS-CoV-2 IgA and IgG in the nasal wash and BALF (Fig 1B-E). Neither IM Prime only nor IN spike only was sufficient for the development of mucosal antibodies. In the serum, IM Prime only was sufficient to induce low levels of IgA and IgG; however, Prime and Spike led to significant systemic boosting of both anti-spike S1 IgA and IgG (Fig 1F,G). These increases in antibody level correlated with increases in neutralization titers both in the BALF and serum (Fig 1H-K). These results indicate that single-dose unadjuvanted intranasal spike alone is not immunogenic, and that induction of a potent mucosal and systemic antibody response by unadjuvanted spike requires prior systemic priming, in this case by mRNA-LNP.

Tissue resident memory B cells (B_RM_) in the lungs have been shown to assist in rapid recall response of antibody secreting B cells upon secondary heterologous challenge in mouse influenza models and may be an important local immune effector in protecting against SARS-CoV-2 (*30*). Using intravenous (IV) CD45 labeling combined with B cell tetramers specific for receptor binding domain (RBD) of the spike protein, we found that Prime and Spike leads to increased antigen specific B cells within lung tissue (IV^-^CD19^+^B220^+^Tetramer^+^) (Fig 1L). Given that the tetramer only assessed for RBD binding, we also looked at the polyclonal tissue response which likely represents a more complete set of B cells reactive to the entire spike within lung tissue. We found increases in class switched antibody secreting cells (ASC) (IV^-^ CD19^+/-^CD138^+^) in lung tissue expressing IgA or IgG (Fig 1M,N), and we found increased class switched B_RM_ (IV^-^CD19^+^B220^+^IgD^-^IgM^-^CD38^+^) expressing IgA or IgG (Fig 1O,P). These results are consistent with increased mucosal antibody production and indicate that Prime and Spike elicits local B cell responses in the lung.

### Prime and Spike induces mucosal T cell immunity

Given that we found that Prime and Spike induced mucosal humoral memory responses in the respiratory tract, we next wanted to assess the induction of lung tissue resident memory T cells (T_RM_). While subunit vaccines have traditionally not been potent inducers of antigen specific T cell responses, we hypothesized that the immune memory generated by mRNA-LNP priming, which has been shown to be sufficient for induction of T cell memory responses in both animal models and humans, would enable subunit mediated T cell boosting responses. Similar to above, we combined CD45 IV labeling to differentiate circulating from immune cells within lung tissue with major histocompatibility complex (MHC) class I tetramer to a conserved sarbecovirus spike epitope (VNFNFNGL). We found significant induction of spike IV^-^ tetramer^+^ CD8^+^ T cells, which expressed canonical markers of T_RM_ including CD69^+^ and CD103^+^, within lung tissue (Fig 2B-D), the lower airway BALF (Fig 2E-G), and in the upper airway nasal turbinate (Fig 2H-J). Additionally, we found significant increases in antigen experienced CD4^+^ T cells (IV^-^ CD44^+^CD4^+^), many of which also expressed markers of T_RM_ CD69^+^ and CD103^+^ both within lung tissue (Fig 2K-M) and from lower airway recovered from BALF (Fig 2N-P). These results indicate that Prime and Spike not only induces humoral mucosal responses, but also robustly elicits lung parenchyma and airway CD8^+^ T_RM_ and CD4^+^ T_RM_.

**Fig 2.**
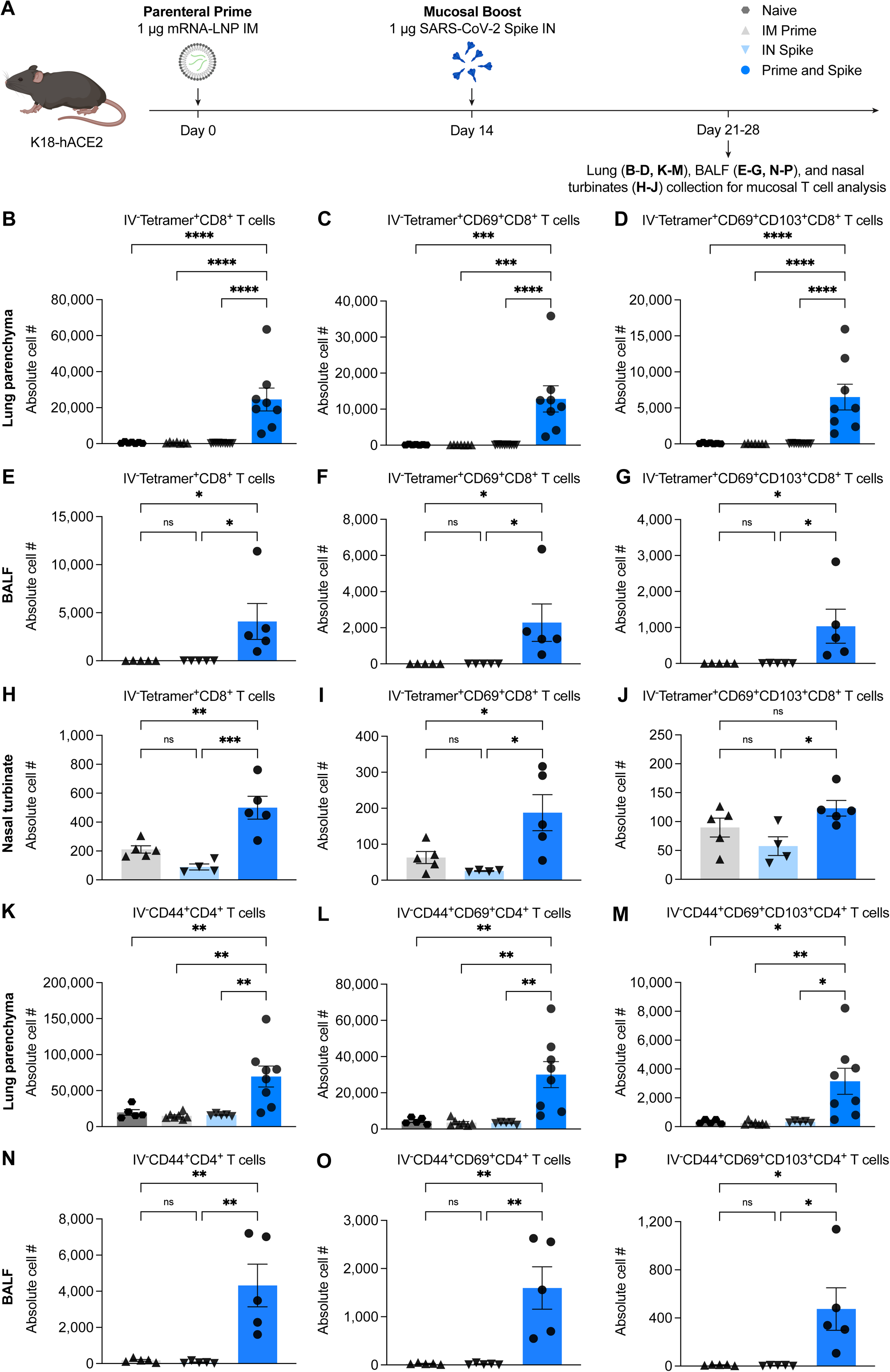
IN boosting with stabilized SARS-CoV-2 spike induces mucosal T cell memory. (**A**) Experimental schema: K18-hACE2 mice were IM primed with 1 μg mRNA-LNP and 14 days later IN boosted with 1 μg SCV2 spike. Lung tissues, BALF, and nasal turbinates were collected for extravascular T cell analysis. Lung tissues were collected 14 days post boost, BALF and nasal turbinates 7 days post boost. (**B-H**) Extravascular CD8 T cell responses: Quantification of SCV2 spike-specific Tetramer^+^ CD8 T cells, CD69^+^CD103^-^Tetramer^+^ CD8 T cells, or CD69^+^CD103^+^Tetramer^+^ CD8 T cells in lung tissues (**B-D**), BALF (**E-G**), or nasal turbinates (**H-J**) from naïve, IM Prime, IN Spike, or Prime and Spike mice. (**K-P**) Extravascular CD4 T cell responses: Quantification of activated polyclonal CD4 T cells, CD69^+^CD103^-^ CD4 T cells, or CD69^+^CD103^+^ CD4 T cells in lung tissues (**K-M**) or BALF (**N-P**) from naïve, IM Prime, IN Spike, or Prime and Spike mice. Mean ± s.e.m.; Statistical significance was calculated by one-way ANOVA followed by Tukey correction (**B-P**); *P ≤ 0.05, **P ≤ 0.01, ***P ≤ 0.001, ****P ≤ 0.0001. Data are representative of three independent experiments.

### Delayed interval Prime and Spike is sufficient to induce mucosal immunity

While we showed that Prime and Spike at a 14-day interval between priming and boosting resulted in significant induction of mucosal humoral and cellular immune memory responses, we wondered whether a delayed boost could also induce significant mucosal humoral and cellular responses. To test this question, K18-hACE2 mice that received 1 μg IM Prime were boosted with IN Spike 84 days later. We sampled humoral and cellular mucosal immune responses at day 91 (7 days post boost) and day 140 (56 days post boost) (Fig S1A). We found that delayed IN Spike was sufficient to induce CD8^+^ T_RM_ which persisted for at least 56 days (Fig S1B-D). CD4^+^ T_RM_ were induced early at 7 days post boost; however, their longevity seemed to wane by 56 days, at least polyclonally (Fig S1E-G). Similar to the CD8^+^ T_RM_ response, we found not only adequate humoral response to delayed boosting, but strong and increasing mucosal IgA and IgG in BALF (Fig S1H,I), and strong and increasing serum IgA and IgG (Fig S5D,E) at 56 days post boosting. These results indicate that Prime and Spike given with a 3-month interval between doses is sufficient to elicit long lasting mucosal and systemic humoral and cellular immune responses.

### IN delivery of mRNA polyplexes also mediates mucosal boosting

We next assessed the ability of alternative platforms for IN Spike boosting. Poly(amine-co-ester)s (PACE) are biodegradable terpolymers that have been developed to encapsulate and deliver nucleic acids such as mRNA or DNA to specified tissues *in vivo* depending on the properties of the polymer (*31*). Recent studies have shown that mRNA-LNP delivered to the respiratory tract is lethal in a dose dependent manner in mice (*32*). In contrast, PACE materials have been developed to be relatively immunologically silent, enabling administration to locations more susceptible to immunopathology such as the respiratory tract. To assess the safety and efficacy of PACE encapsulating mRNA encoding spike protein, mRNA was extracted from Comirnaty and encapsulated in PACE polyplexes. For vaccination, K18-hACE2 mice were injected with 1 μg IM Prime (mRNA-LNP), and 14 days later received 1 μg of Mrna encapsulated in PACE and administered IN (PACE-Spike). Additional control groups included PACE-Spike only and IM Prime + extracted mRNA without PACE encapsulation (naked mRNA) (Fig 3A). Similar to what we found with Prime and Spike, Prime and PACE-Spike induced antigen specific CD8^+^ T_RM_ (IV^-^Tetramer^+^) expressing canonical tissue residency markers (CD69^+^ and CD103^+^) (Fig 3B-D). Additionally, PACE-Spike boosted mice developed high levels of BALF anti-SARS-CoV-2 IgA; levels of BALF IgG and serum IgA and IgG were similar to IM Prime only mice (Fig 3E-H). IM Prime followed by IN naked mRNA was unable to induce mucosal or systemic immune responses above that of IM Prime alone indicating that mRNA encapsulation by PACE was required for mucosal boosting. Additionally, a single dose of IN PACE-Spike alone was insufficient to elicit any detectable mucosal or systemic antibody response at this dose.

**Fig 3.**
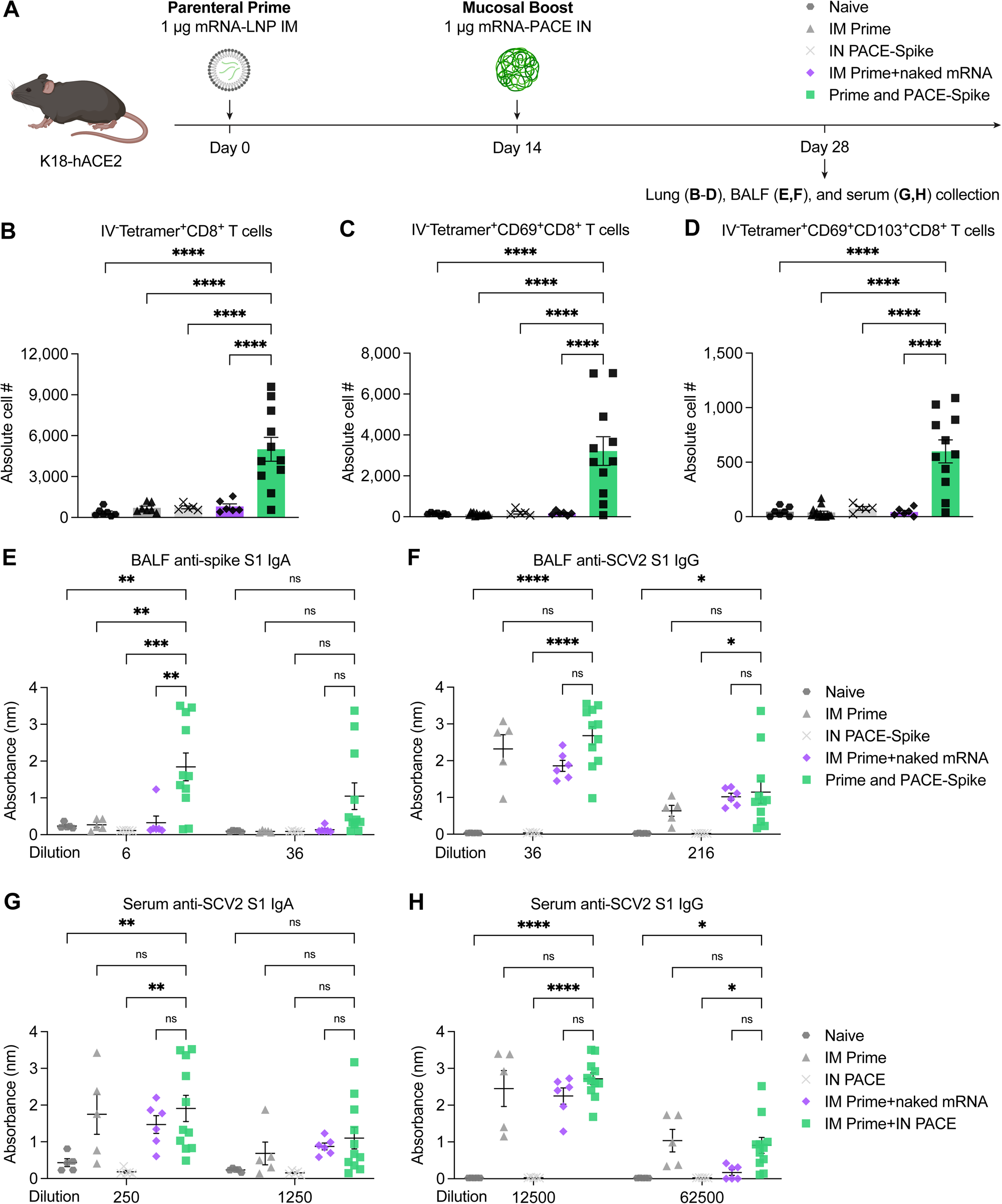
IN delivery of SCV2 spike mRNA encapsulated in poly(amine-co-ester) (PACE) terpolymers mediates mucosal boosting. (**A**) Experimental schema: K18-hACE2 mice were IM primed with 1 μg of mRNA-LNP, followed by IN boosting with 1 μg of naked mRNA (IN naked mRNA) or 1 μg of mRNA encapsulated by PACE (IN PACE-Spike) 14 days post IM Prime. Fourteen (14) days post IN boost, BALF and blood were collected for antibody measurement. Lung tissues were collected for CD8 T cell analysis. (**B-D**) Quantification of total Tetramer^+^ CD8 T cells, CD69^+^CD103^-^Tetramer^+^ CD8 T cells, or CD69^+^CD103^+^Tetramer^+^ CD8 T cells in lung tissues from naïve, IM Prime, IN PACE-Spike, IM Prime+IN naked mRNA, or Prime and PACE-Spike mice. (**E-H**) Measurement of SARS-CoV-2 spike S1 subunit-specific BALF IgA (**E**), BALF IgG (**F**), serum IgA (**G**), and serum IgG (**H**) in naïve, IM Prime, IN PACE-Spike, IM Prime+IN naked mRNA, or Prime and PACE-Spike mice. Mean ± s.e.m.; Statistical significance was calculated by one-way ANOVA followed by Tukey correction (**B-D**) or two-way ANOVA followed by Tukey correction (**E-H**); *P ≤ 0.05, **P ≤ 0.01, ***P ≤ 0.001, ****P ≤ 0.0001. Data are pooled from two independent experiments.

### Prime and Spike or Prime and PACE-Spike in the context of waning mRNA-LNP immunity protects against lethal SARS-CoV-2 challenge

While current vaccines were initially extremely effective at eliciting protective immunity, waning antibody levels and immune evasion will necessitate boosters against SARS-CoV-2 for the foreseeable future; however, the best method for boosting remains a question. To test whether IN administration would provide an alternative protective boost, we utilized a low dose mRNA-LNP vaccine challenge model to mimic waning immunity; we performed single dose immunization with 0.05 μg mRNA-LNP. While these low dose mRNA-LNP vaccinated mice uniformly develop systemic antibody responses, we have previously shown that this dose is not sufficient to protect from SARS-CoV-2 challenge (*15*). Fourteen days post prime mice received IN Spike (1 μg unadjuvanted spike protein). Similar to 1 μg IM Prime mice that we described earlier, 0.05 μg IM primed mice boosted with IN Spike developed a significant increase in antigen specific CD8^+^ T_RM_ in the lungs as well as IgA and IgG in the BALF at 42 days post boost (Fig. S2). These data also indicate that even very low levels of immune memory generated by low dose mRNA-LNP prime can be effectively boosted to induce mucosal and systemic humoral and cellular memory by unadjuvanted IN spike.

Naïve, low dose Prime only, and low dose Prime and Spike mice were then challenged with 6×10^4^ PFU homologous/ancestral WA1 strain SARS-CoV-2. Mice were either euthanized at 2 DPI and viral burden assessed by plaque assay from nasal turbinates and lungs, euthanized at 5 DPI and lungs assessed for pathology, or monitored for weight loss and mortality for 14 days (Fig. 4A). All mice given Prime and Spike were completely protected from weight loss or death, but neither naïve nor low dose Prime only mice were protected from viral challenge (Fig 4B-D). Additionally, this significant improvement in morbidity and mortality in mice receiving Prime and Spike was accompanied by reduced viral burden in both the upper respiratory tract (nasal turbinates) and lower respiratory tract (lungs) (Fig 4E,F). Further, Prime and Spike led to significant protection from lung pathology with only 1 of 6 mice developing limited mononuclear infiltrates at 5 DPI, while the remaining mice were completely protected with lung architecture similar to that seen in uninfected mice (Fig, 4G,H). To assess the protective capacity of a mRNA-PACE IN boost, we again made use of the low-dose prime mRNA-LNP mice, and boosted with 10 μg of mRNA-PACE IN. We found that Prime and PACE-Spike resulted in significant protection from morbidity and mortality (Fig 4I-L). These data suggest that either IN unadjuvanted spike or mRNA-PACE encoding spike sufficiently boost mucosal immunity to protect from COVID-19 like pulmonary disease and mortality in a preclinical mouse model. These results also highlight the robustness, versatility, and safety of this vaccine strategy as intranasal boosting of systemic mRNA-LNP priming by either modality is sufficient to induce mucosal immunity and to provide protection against lethal SARS-CoV-2 challenge.

**Fig 4.**
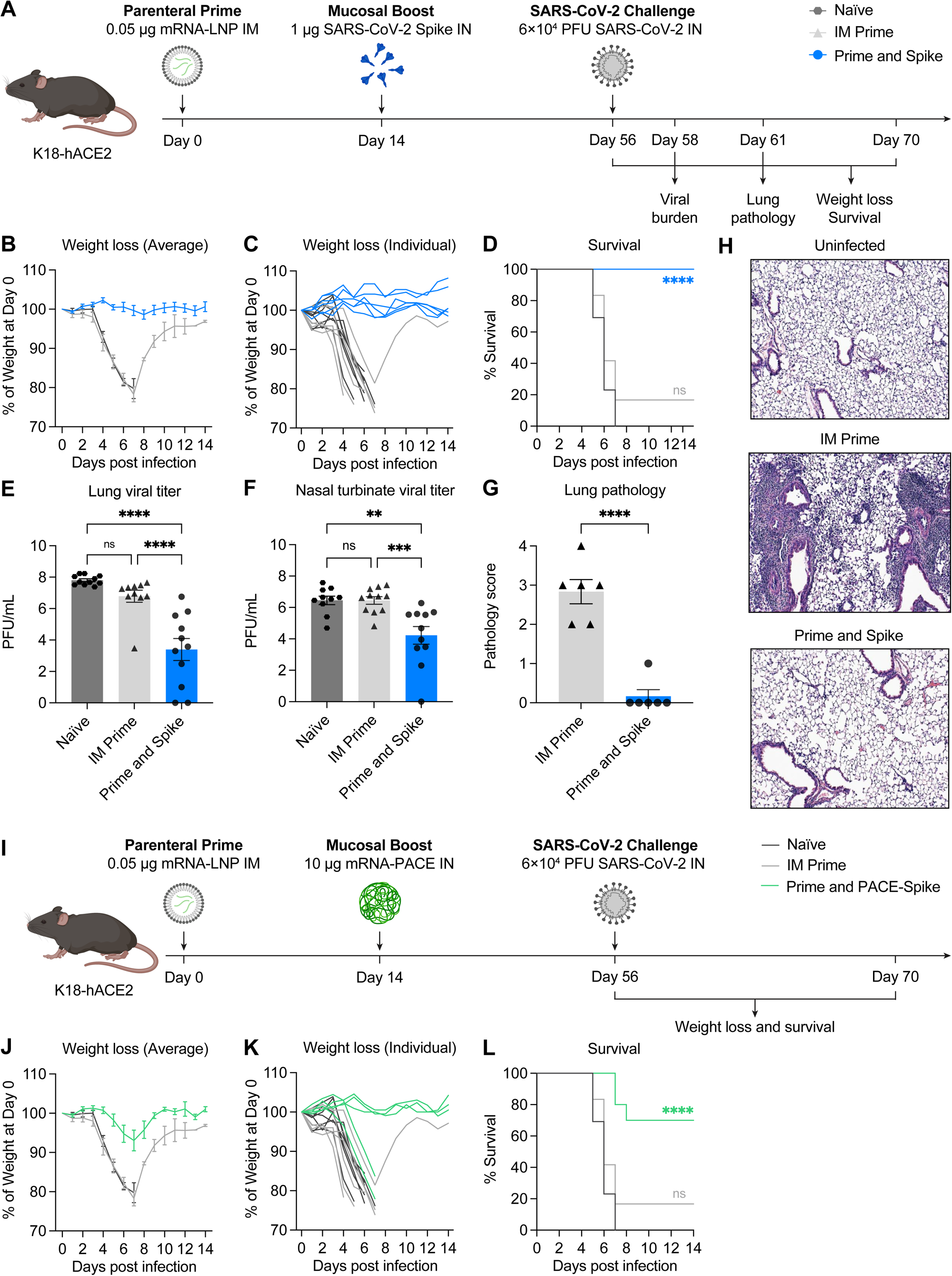
Intranasal stabilized SARS-CoV-2 spike boosting protects against COVID-19-like disease. (**A**) Experimental schema: K18-hACE2 mice were IM primed with 0.05 μg of mRNA-LNP and IN boosted with 1 μg of spike IN 14 days post IM Prime. 6 weeks post boost, mice were challenged with 6 × 10^4^ PFU SARS-CoV-2 (2019n-CoV/USA_WA1/2020). The first cohort was used to evaluate weight loss and survival up to 14 days post infection (DPI). The second cohort was used to collect lung and nasal turbinate tissues 2 DPI for viral titer measurement. The third cohort was used to collect lung tissues 5 DPI for histological assessment. (**B-D**) Weight loss and survival of naïve, IM Prime, or Prime and Spike mice from 1 to 14 DPI. (**E-F**) Measurement of infectious virus titer in lung and nasal turbinate tissues at 2 DPI by plaque assay. (**G**) Pathology score of lung sections at 5 DPI by Hematoxylin and Eosin (H&E) staining. (**B**) Representative H&E staining results from uninfected, IM Prime, or Prime and Spike mice. (**I**) Experimental schema: K18-hACE2 mice were IM primed with 0.05 μg of mRNA-LNP and IN boosted with 10 μg of mRNA encapsulated by PACE (IN PACE-Spike) 14 days post IM Prime. 6 weeks post boost, mice were challenged with 6 × 10^4^ PFU SARS-CoV-2 (2019n-CoV/USA_WA1/2020). Weight loss and survival were monitored up to 14 DPI. (**J-L**) Weight loss and survival of naïve, IM Prime, or Prime and PACE-Spike K18-hACE2 mice from 1 to 14 DPI. Mean ± s.e.m.; Statistical significance was calculated by log-rank Mantel–Cox test (**D**,**L**), one-way ANOVA followed by Tukey correction (**E**,**F**), or student’s t-test (**G**); *P ≤ 0.05, **P ≤ 0.01, ***P ≤ 0.001, ****P ≤ 0.0001. Data are pooled from two independent experiments.

## Prime and Spike elicits robust systemic immunity similar to parenteral mRNA-LNP based boost

IM injected mRNA-LNP based vaccines are the current standard recommended boosting strategy in many countries as immunogenicity and vaccine efficacy studies have most concentrated on this method of boosting. To compare Prime and Spike to IM mRNA-LNP prime/boost, we primed K18-hACE2 mice with 1 μg of mRNA-LNP, followed 14 days later by either 1 μg IN Spike or 1 μg IM mRNA-LNP. Mice were euthanized 31 days post boost and antigen specific CD8^+^ T_RM_ were assessed by flow cytometry, antibodies from BALF and serum were assessed by ELISA, and VSV pseudovirus neutralization assay was performed to assess serum antibody neutralization response (Fig S3A). We found that both IM mRNA-LNP boosted and IN Spike boosted animals developed increased levels of extravascular (IV^-^) Tetramer^+^CD8^+^ T cells; however, only IN spike boosted animals developed CD8^+^ T_RM_ that express CD69^+^ and CD103^+^ in the lung (Fig 3SB-D). By ELISA we found that only IN spike boosted animals developed anti-SARS-CoV-2 IgA in the BALF. BALF IgG levels were similar in IM mRNA-LNP and IN spike boosted mice, possibly representing transcytosis from elevated systemic antibody levels in the IM mRNA-LNP boosted mice. We similarly found equivalent serum levels of anti-SARS-CoV-2 IgA and IgG in IM mRNA-LNP and IN spike boosted mice. Neutralization assays from serum also showed similar IC50 between IM mRNA-LNP and IN Spike boosting. These data demonstrate that Prime and Spike induces similar systemic neutralizing antibody levels to IM mRNA-LNP boosting, which has been shown to be a correlate of protection, and uniquely elicits mucosal IgA and CD8^+^ T_RM_.

### Heterologous spike robustly elicits cross-reactive immunity without original antigenic sin

The experiments above clearly demonstrate that boosting at a distinct anatomic location, in this case the respiratory mucosa, either by unadjuvanted subunit spike or by PACE-Spike encoding spike, enables the formation of new mucosal immune memory at the newly boosted site and enhances systemic immunity to that antigen. In both unadjuvanted subunit spike and mRNA-PACE, the boosting antigen is homologous to the systemic priming antigen (mRNA-LNP). Current circulating strains of SARS-CoV-2, notably Delta and Omicron, have significant changes to the spike protein sequence and structure. Delta harbors T19R, G142D, Δ156-157, R158G, Δ213-214, L452R, T478K, D614G, P681R, and D950N mutations, whereas Omicron harbors A67V, Δ69-70, T95I, G142D, Δ143-145, N211I, L212V, ins213-214RE, V215P, R216E, G339D, S371L, S373P, S375F, K417N, N440K, G446S, S477N, T478K, E484A, Q493R, G496S, Q498R, N501Y, Y505H, T547K, D614G, H655Y, N679K, P681H, N764K, D796Y, N856K, Q954H, N969K, and L981F mutations. These mutations have made both Delta and Omicron transmit more rapidly and evade pre-existing humoral immunity, and it is likely that future variants will diverge even more, suggesting a boosting strategy that elicits broadly reactive immunity will be necessary to neutralize future variants. To test the ability of an unadjuvanted heterologous spike protein for IN boosting, we primed K18-hACE2 mice with 1 μg mRNA-LNP followed 14 days later by boosting with 5 μg of SARS-CoV-1 spike containing trimer stabilizing mutations (R667A, K968P, V969P), or Prime and SpikeX (Fig 5A). While SARS-CoV-1 is a related sarbecovirus, its spike protein only shares 76% homology with the original SARS-CoV-2 spike sequence that is encoded by currently used mRNA-LNP vaccines. For comparison we also boosted mRNA-LNP primed mice with 1 μg IM mRNA-LNP. At 31 days post boost, using CD45 IV labeling, we found significantly increased IV^-^ Tetramer^+^ CD8^+^ T cells that express canonical T_RM_ markers CD69^+^ and CD103^+^ (Fig 5B-D). As noted earlier, this MHC I tetramer sequence is highly conserved within the sarbecovirus family, which both SARS-CoV-1 and SARS-CoV-2 are a part of. Next, we assessed the development of anti-SARS-CoV-1 antibodies in BALF and serum and found significant increases of anti-SARS-CoV-1 IgA and IgG in both the respiratory mucosa and the circulation in Prime and SpikeX relative to IM mRNA-LNP prime/boost. Consistent with previous studies, we find that two doses of SARS-CoV-2 mRNA-LNP is sufficient to induce detectable antibodies that bind SARS-CoV-1 spike (*33*). Next, we assessed anti-SARS-CoV-2 antibodies in BALF and serum. We found that Prime and SpikeX induced higher BALF IgA than IM SARS-CoV-2 mRNA-LNP prime/boost. We found similar levels of anti-SARS-CoV-2 IgG in BALF which likely represents the increased serum IgG that we found (Fig 5I-L). Next, using VSV-based pseudovirus neutralization assay we show that serum from Prime and SpikeX mice develops higher neutralization titers against SARS-CoV-1 than mice boosted with IM SARS-CoV-2 mRNA-LNP (Fig 5M,N). Similarly and consistent with serum IgG levels, IM SARS-CoV-2 mRNA-LNP prime/boost mice have significantly higher neutralization titers against SARS-CoV-2 than Prime and SpikeX mice (Fig 5O,P). Taken together, these data indicate that IN boosting with unadjuvanted heterologous spike protein can induce potent mucosal cellular and humoral memory against significantly divergent spike protein in the absence of original antigenic sin.

**Fig 5.**
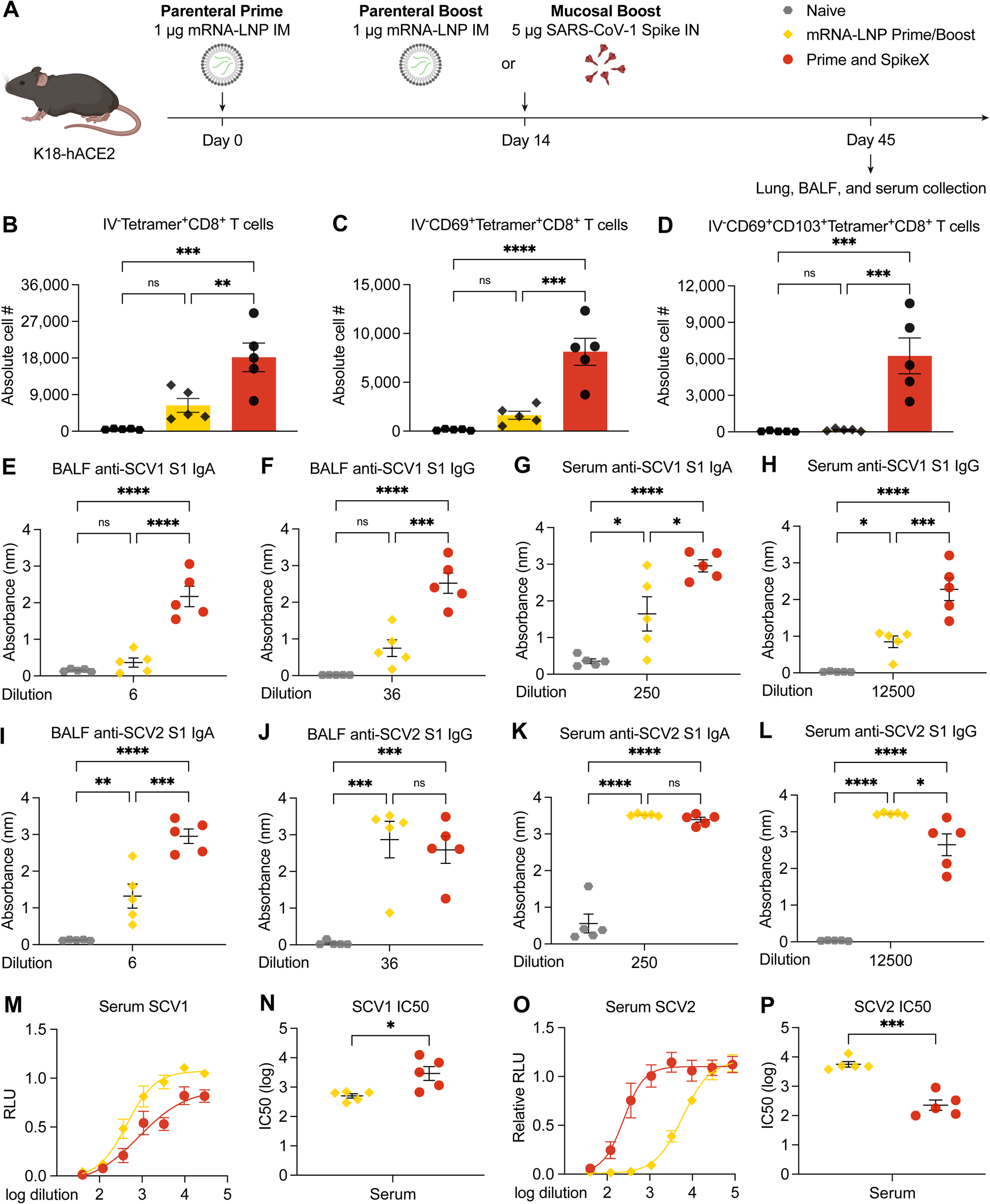
Heterologous IN boosting with SARS-CoV-1 spike enhances pre-existing SCV2-specific immunity and broadens reactivities to SCV1. (**A**) Experimental schema: K18-hACE2 mice were IM primed with 1 μg of mRNA-LNP, followed by boosting with 1 μg of mRNA-LNP IM, or 5 μg of prefusion-stabilized, trimeric, recombinant SARS-CoV-1 (SCV1) spike IN (IN SpikeX) 14 days post IM Prime. 31 days post boost, lung tissues were collected for T cell analysis by flow cytometry, and BALF and blood were collected for antibody measurement. (**B-D**) Quantification of total Tetramer^+^ CD8 T cells, CD69^+^CD103^-^Tetramer^+^ CD8 T cells, or CD69^+^CD103^+^Tetramer^+^ CD8 T cells in lung tissues from naïve, mRNA-LNP Prime/Boost, or Prime and SpikeX mice. (**E-H**) Measurement of SCV1 spike S1 subunit-specific BALF IgA (**E**), BALF IgG (**F**), serum IgA (**G**), and serum IgG (**H**) in naïve, mRNA-LNP Prime/Boost, or Prime and SpikeX mice. (**I-L**) Measurement of SCV2 spike S1 subunit-specific BALF IgA (**I**), BALF IgG (**J**), serum IgA (**K**), and serum IgG (**L**) in naïve, mRNA-LNP Prime/Boost, or Prime and SpikeX mice. (**M**,**N**) Measurement of neutralization titer against SCV1 spike-pseudotyped VSV. (**O**,**P**) Measurement of neutralization titer against SCV2 spike-pseudotyped VSV. Mean ± s.e.m.; Statistical significance was calculated one-way ANOVA followed by Tukey correction (**B-L**) or student’s t-test (**N**,**P**); *P ≤ 0.05, **P ≤ 0.01, ***P ≤ 0.001, ****P ≤ 0.0001.

## Discussion

Here we describe the preclinical development of a new vaccine strategy, Prime and Spike, where IN unadjuvanted spike subunit protein can elicit robust protective mucosal immunity following mRNA-LNP parenteral immunization. Prime and Spike elicited robust mucosal cellular and humoral memory responses including the establishment of CD8^+^ T_RM,_ CD4^+^ T_RM,_ B_RM,_ IgA, and IgG. We found that an IN unadjuvanted spike booster can be administered months out from primary immunization, and that it offers comparable systemic neutralizing antibody responses to mRNA-LNP boost, with the added benefit of mucosal immunity. Similarly, we find that IN boosting with mRNA encapsulated in PACE polymers (Prime and PACE-Spike) elicits increased mucosal CD8^+^ T_RM_ and IgA. Both boosting methods resulted in protection from lethal SARS-CoV-2 challenge in a mouse model of waning immunity. Finally, by utilizing a divergent spike antigen, we demonstrate that Prime and SpikeX can generate mucosal immunity to SARS-CoV-1, while also boosting the neutralizing antibodies to the original antigenic target, SARS-CoV-2 spike.

While the goal of vaccination has been to prevent individual morbidity and mortality, the evolution of SARS-CoV-2 throughout the pandemic has highlighted the need for rapidly deployable mucosal vaccines which not only protect the individual, but also prevent transmission. mRNA-LNP based vaccines initially showed incredible efficacy (∼95%) against severe disease as well as reduced transmission; however, waning immunity, viral immune evasion, and increased viral transmissibility have demonstrated their limits in current form.

Specifically, these and other currently approved SARS-CoV-2 vaccines induce little mucosal immunity in the respiratory tract (*15-17*), the site of infection and transmission. Our data demonstrated that Prime and Spike significantly reduced the viral load in the nasal cavity and the lung compared to parenteral vaccine alone, indicating the promise of Prime and Spike in reducing infection and transmission. Improving upon current vaccine platforms to provide mucosal immunity is important to curb this current pandemic, and certainly will be important to combat the next.

Preclinical studies of both SARS-CoV-2 and Influenza have demonstrated that intranasal vaccination decreases viral shedding and transmission relative to parenteral vaccines (*19-23*). Despite these studies, there is only one currently approved respiratory mucosal vaccine, Flumist, which relies on a live cold-adapted influenza virus. While this is an effective and well-studied technology, it is contraindicated in people with underlying respiratory conditions, is only approved for the young, and is not amenable to rapid implementation in the setting of an emergent respiratory virus epidemic or pandemic as they require extensive research and development. As such, all current early phase clinical trials of mucosal administered SARS-CoV-2 vaccines rely on either replication deficient or attenuated viral vectors; however, safety and efficacy are yet to be established, especially given that preexisting immunity to these vectors can lead to reduced immunogenicity (*34*). In fact, some vector based mucosal vaccines—including two Merck candidates V590 and V591—have already been abandoned after phase 1 clinical trials showed poor immunogenicity.

This is the first report of boosting preexisting systemic immunity generated by currently approved SARS-CoV-2 vaccines (mRNA-LNP) with IN unadjuvanted spike protein to elicit robust mucosal immunity. This technology enables the use of a non-inflammatory local boost at a tissue compartment sensitive to immunological effector mechanisms such as the respiratory tract to induce mucosal and systemic immunity. We also demonstrate that IN unadjuvanted subunit boost induces CD4^+^ T_RM_, CD8^+^ T_RM_, B_RM_, and mucosal IgA and IgG, and challenges prior dogma suggesting that subunit vaccines cannot induce robust cellular responses and that adjuvants are required for subunit vaccines to induce respiratory mucosal immunity. Further, this is the first report to successfully and safely deliver mRNA to the respiratory tract as a SARS-CoV-2 vaccine boost utilizing a novel formulation of biodegradable, non-inflammatory PACE polymers.

These technologies are likely broadly applicable to be used as boosters against new VOCs for SARS-CoV-2 or as part of a primary immunization strategy for newly emerging respiratory pathogens. While it is possible that these results rely on specific characteristics of the mRNA-LNP priming, we believe that this will likely work with other primary immunization regimens, or in the case of previous infection; however, this will require further assessment. Additionally, it has been shown that the highly stabilized spike enhances its immunogenicity, and that applying this novel vaccination strategy to other pathogens may require the addition of stabilizing mutation of the antigen of choice to enable unadjuvanted boosting as reported here. That said, using a less modified SARS-CoV-1 spike also enabled unadjuvanted IN boosting of a heterologous antigen.

Vaccines that generate broadly neutralizing serum immunity against a wide variety of sarbecoviruses has been a recent goal to combat both newly emerging SARS-CoV-2 variants and other potential future pandemic SARS like coronaviruses. Here, we utilize SARS-CoV-1 spike as a heterologous IN boost, in Prime and SpikeX, and show that prior SARS-CoV-2 mRNA-LNP does not prevent the development of SARS-CoV-1 neutralizing antibodies, but that it likely enables it as IN Spike alone was not immunogenic. Prime and SpikeX simultaneously elicits more broadly reactive neutralizing antibodies and mucosal immunity. While some recent studies have successfully reported the development of pan-sarbecovirus vaccines (*33, 35*), this is the first report that we are aware of that shows the induction of both systemic and mucosal immunity against both SARS-CoV-1 and SARS-CoV-2.

SARS-CoV-2 will continue to evolve and become more immune evasive and transmissible as we have seen with the Omicron VOC. We will be requiring boosting in human populations for the foreseeable future; however, boosting that induces mucosal immunity may help to enhance protection and slow transmission as these new variants emerge.

## Materials and Methods

All procedures were performed in a BSL-3 facility (for SARS-CoV-2–infected mice) with approval from the Yale Institutional Animal Care and Use Committee and Yale Environmental Health and Safety.

### Cell and virus

As reported in previous manuscripts (*15, 36, 37*), Vero E6 cells over expressing hACE2 and TMPRSS2 (kindly provided by Barney Graham NIH-VRC) were cultured in Dulbecco’s Modified Eagle Medium (DMEM) supplemented with 1% sodium pyruvate and 5% fetal bovine serum (FBS) at 37°C and 5% CO_2_. SARS-CoV-2 isolate hCOV-19/USA-WA1/2020 (NR-52281) was obtained from BEI Resources and was amplified in VeroE6 cells over expressing hACE2 and TMPRSS2. Cells were infected at a MOI 0.01 for two-three days to generate a working stock and after incubation the supernatant was clarified by centrifugation (500 g × 5 min) and filtered through a 0.45-micron filter and stored at -80°C. Viral titers were measured by standard plaque assay using Vero E6 cells over expressing hACE2 and TMPRSS2.

### Mice

B6.Cg-Tg(K18-ACE2)2Prlmn/J (K18-hACE2) mice were purchased from The Jackson Laboratory and subsequently bred and housed at Yale University. Eight to twelve-week-old female were used for immunization experiments. All procedures used in this study (sex-matched, age-matched) complied with federal guidelines and the institutional policies of the Yale School of Medicine Animal Care and Use Committee.

### SARS-CoV-2 infection

Mice were anesthetized using 30% v/v Isoflurane diluted in propylene glycol. Using a pipette, 50 μL containing 6×10^5^ PFU SARS-CoV-2 was delivered intranasally.

### mRNA Extraction from Comirnaty mRNA-LNP

mRNA was extracted from the vaccine formulation with a TRIzol/chloroform separation method described here (*38*). Briefly, aliquots of vaccine were dissolved in TRIzol LS (Thermo Fisher Scientific) at 1:6.6 vaccine to TRIzol volume ratio. Following a 15 min incubation (37°C, shaking) 0.2 mL of chloroform was added per 1 mL of TRIzol. The solution was shaken vigorously for 1 min and then incubated at room temperature for 3 min. The solution was centrifuged at 12,000 x g for 8 min at 4°C. The aqueous layer containing the isolated mRNA was further purified with a RNeasy Maxi Kit purchased from Qiagen (Germantown, MD, USA) following the manufacturers protocol. The RNA was eluted from the column on the final step with sodium acetate buffer (25 mM, pH 5.8) warmed to 37°C. Extracted mRNA was analyzed for concentration and purity by NanoDrop measurements of the absorbance at 260, 280 and 230 nm, with purity being assessed as A260/A280 > 2 and A260/A230 > 2. Agarose gel electrophoresis was used to determine the length and verify that the mRNA remained intact. Extracted mRNA containing 1:100 SYBR Safe stain (Thermo Fisher Scientific) was loaded onto a 1% agarose gel and run at 75V with TAE buffer containing 1:5000 SYBR Safe stain (Fig S4).

### PACE Polyplex Formulation and Characterization

PACE polymers were synthesized and characterized as previously described (*39*). All polyplexes were formulated at a 50:1 weight ratio of polymer to mRNA. PACE polymers were dissolved at 100 mg/mL overnight in DMSO (37°C, shaking). Prior to polyplex fabrication, an optimal PACE polymer blend was produced by mixing solutions of PACE polymers containing an end-group modification (*40*) and a polyethylene glycol tail (*31*). mRNA and polymer were diluted into equal volumes of sodium acetate buffer (25 mM, pH 5.8). The polymer dilution was then vortexed for 15 s, mixed with the mRNA dilution, and vortexed for an additional 25 s. Polyplexes were incubated at room temperature for 10 min before use.

### Vaccination

Used vials of Comirnaty vaccine were acquired from Yale Health pharmacy within 24 hr of opening and stored at 4°C. Vials contained residual vaccine (diluted to 100 μg/mL per manufacturer’s instructions) which was removed with spinal syringe and pooled. Pooled residual vaccine was aliquoted and stored at -80°C. Mice were anaesthetized using a mixture of ketamine (50 mg/kg) and xylazine (5 mg/kg), injected intraperitoneally. Vaccine was diluted in sterile PBS and 10 μL or 20 μL was injected into the left quadriceps muscle with a 31 g syringe for a final dose of 1 μg or 0.05 μg as indicated. For intranasal vaccination with SARS-CoV-2 stabilized spike (ACRO biosystems, SPN-C52H9) or SARS-CoV-1 spike (ACRO biosystems, SPN-S52H6) was reconstituted in sterile endotoxin free water according to the manufacturers protocol, and then diluted in sterile PBS and stored at -80°C. Mice were anesthetized using 30% v/v Isoflurane diluted in propylene glycol and administered 1 μg or 5 μg (as indicated) in 50 μL via the IN route. For IN mRNA-PACE, 50 μL of polyplexes in solution was given at the indicated dose.

### Viral titer analysis

At indicated time points mice were euthanized in 100% Isoflurane. ∼50% of total lung was placed in a bead homogenizer tube with 1 mL of PBS with 2% FBS and 2% antibiotics/antimycotics (Gibco) and stored at -80°C. Lung homogenates were cleared of debris by centrifugation (3900 rpm for 10 min). Infectious titers of SARS-CoV-2 were determined by plaque assay in VeroE6 cells over expressing hACE2 and TMPRSS2 in DMEM supplemented with NaHCO_3_, 2% FBS, and 0.6% Avicel RC-581. Plaques were resolved at 40-42 hours post infection by fixing in 10% Neutral Buffered Formalin for 1 hour followed by staining for 1 hour in 0.5% crystal violet in 20% ethanol for 30 min. Plates were rinsed in water to visualize plaques.

### SARS-CoV-2 specific-antibody measurements

ELISAs were performed as previously described (*36, 41*) with modifications noted and reproduced here for convenience. 96-well MaxiSorp plates (Thermo Scientific #442404) were coated with 50 μL/well of recombinant SARS-CoV-2 S1 protein (ACRO Biosystems S1N-C52H3) or SARS-CoV-1 S1 protein (ACRO Biosystems S1N-S52H5) at a concentration of 2 μg/mL in PBS and were incubated overnight at 4°C. The coating buffer was removed, and plates were incubated for 1 hour at RT with 250 μL of blocking solution (PBS with 0.1% Tween-20, 5% milk powder). Serum or bronchoalveolar lavage fluid (BALF) was diluted in dilution solution (PBS with 0.1% Tween-20 and 2% milk powder) and 100 μL of diluted serum or BALF was added and incubated for two hours at RT. Plates were washed five times with PBS-T (PBS with 0.05% Tween-20) with automatic plate washer (250 μL per cycle) and 50 μL of HRP anti-mouse IgG (Cell Signaling Technology #7076, 1:3,000) or HRP anti-mouse IgA (Southern Biotech #1040-05, 1:1,000) diluted in dilution solution added to each well. After 1 h of incubation at RT, plates were washed three times with PBS-T in automatic plate washer. Plates were developed with 50 μL of TMB Substrate Reagent Set (BD Biosciences #555214) and the reaction was stopped after 15 min by the addition of 50 μL 2 N sulfuric acid. Plates were then read at a wavelength of 450 nm and 570nm, and the difference reported.

### Immunohistochemistry and Pathological analysis

Yale pathology performed embedding, sectioning and H&E staining of lung tissue. A pulmonary pathologist reviewed the slides blinded and identified immune cell infiltration and other related pathologies. Scoring 1-4 as follows: (1) Mild patchy mononuclear infiltrate, parenchymal and perivascular, with variably reactive pneumocytes and stromal rection; (2) Moderate patchy mononuclear infiltrate, parenchymal and perivascular, with variably reactive pneumocytes and stromal rection; (3) Mild, dense mixed infiltrate including mononuclear cells and granulocytes / neutrophils; (4) Moderate, dense mixed infiltrate including mononuclear cells and granulocytes / neutrophils.

### Intravascular labeling, cell isolation, and flow cytometry

To discriminate intravascular from extravascular cells, mice were anesthetized with 30% Isoflurane and injected i.v. with APC/Fire 750 CD45 Ab (30-F11, AB_2572116, BioLegend, #103154) and after 3 min labeling, mice were euthanized. Tissues were harvested and analyzed as previously described (*42*) In short lungs were minced with scissors and incubated in a digestion cocktail containing 1 mg/mL collagenase A (Roche) and 30 μg/mL DNase I (Sigma-Aldrich) in RPMI at 37°C for 45 min. Tissue was then filtered through a 70-μm filter. Cells were treated with ammonium-chloride-potassium buffer and resuspended in PBS with 1% BSA.

Single cell suspensions were incubated at 4°C with Fc block and Aqua cell viability dye for 20 min. Cells were washed once with PBS before surface staining. For T cell analysis, cells were stained with anti-CD103 (BV421, 2E7, AB_2562901, BioLegend #121422), anti-CD3 (BV605, 17A2, AB_2562039, BioLegend #100237), anti-CD44 (BV711, IM7, AB_2564214, BioLegend #103057), anti-CD62L (FITC, MEL-14, AB_313093, BioLegend #104406), anti-CD8a (PerCP/Cy5.5, 16-10A1, AB_2566491, BioLegend #305232), anti-CD69 (PE/Cy7, H1.2F3, AB_493564, BioLegend #104512), anti-CD183 (CXCR3) (APC, CXCR3-173, AB_1088993, BioLegend #126512), anti-CD4 (AF700, GK 1.5, AB_493699, BioLegend #100430), and PE-SARS-CoV-2 S 539-546 MHC class I tetramer (H-2K(b)) for 30 min at 4°C. For B cell analysis, cells were stained with ant-GL7 (Pacific Blue, GL7, AB_2563292, BioLegend #144614), anti-IgM (BV605, RMM-1, AB_2563358, BioLegend #406523), anti-CD138 (BV711, 281-2, AB_2562571, BioLegend #142519), anti-CD19 (BV785, 6D5, AB_11218994, BioLegend #115543), anti-IgA (FITC, polyclonal, AB_2794370, SouthernBiotech #1040-02), anti-B220 (PerCP/Cy5.5, RA3-6B2, AB_893354, BioLegend #103236), PE-SARS-CoV-2 RBD tetramer, anti-CD38 (PE/Cy7, 90, AB_2275531, BioLegend #102718), APC-SARS-CoV-2 RBD tetramer, and anti-IgD (AF700, 11-26c.2a, AB_2563341, BioLegend #405730) for 30 min at 4°C. After washing with PBS, cells were fixed using 4% paraformaldehyde. Cell population data were acquired on an Attune NxT Flow Cytometer and analyzed using FlowJo Software (10.5.3; Tree Star). See (Fig S5) for gating strategy.

### SARS-CoV-2 receptor binding domain B cell tetramer production

Recombinant SARS-CoV-2 Spike RBD His Biotin Protein, CF (R&D/BT10500-050) was incubated at a 4:1 molar ratio with either streptavidin-PE (Prozyme PJRS25) or streptavidin-APC (Prozyme PJ27S) for 30 min at 4°C. Mixture was then purified and concentrated in an Amicon Ultra (50 kDA MWCO) spin column and washed 1x with sterile cold PBS. Concentration was determined on a nanodrop and using fluorophore specific absorbance and tetramers were diluted to 1.0 μM in PBS and stored at 4°C.

### Pseudovirus production

VSV-based pseudotyped viruses were produced as previously described (*15, 43-45*). Vector pCAGGS containing the SARS-CoV-2 Wuhan-Hu-1 spike glycoprotein gene was produced under HHSN272201400008C and obtained through BEI Resources (NR-52310). The sequence of the Wuhan-Hu-1 isolate spike glycoprotein is identical to that of the USA-WA1/2020 isolate. SARS-CoV-1 Spike encoding plasmid was kindly provided by Dr. Vincent Munster and previously described (*46*). 293T cells were transfected with either spike plasmid, followed by inoculation with replication-deficient VSV-expressing Renilla luciferase for 1 hour at 37°C. The virus inoculum was then removed, and cells were washed three times with warmed PBS. The supernatant containing pseudovirus was collected 24 and 48 hours after inoculation, clarified by centrifugation, concentrated with Amicon Ultra centrifugal filter units (100 kDa), and stored in aliquots at -80°C. Pseudoviruses were titrated in Huh7.5 cells to achieve a relative light unit signal of ∼600 times the cell-only control background.

### Pseudovirus neutralization assay

VeroE6 overexpressing hACE2 and TMPRSS2 (Fig. 1) or Huh7.5 cell (Fig 5 and S3) were plated (3×10^4^) in each well of a 96-well plate the day before infection. On the day of infection, serum and BALF were heat-inactivated for 30 min at 56°C. Figure 1 sera were tested at a starting dilution of 1:50 and BALF samples were tested at a starting dilution of 1:4, both with 8 twofold serial dilutions. Figure 5 and S3 sera were tested at a starting dilution of 1:40 with 8 threefold serial dilutions. Serial dilutions mixed 1:1 with pseudoviruses and incubated for 1 hour at 37°C. Growth medium was then aspirated from the cells and replaced with 100 μL of serum/virus mixture. At 24 hours infection media was removed and plates flash frozen at -80°C. 30 μg passive lysis buffer (Promega) was added to each well and plates were incubated for 15 min at RT. 30 μg of Renilla-Glo Luciferase Assay System substrate (Promega) was added to each well and incubated at RT for 15 min. Luminescence was measured on a microplate reader (SpectraMax i3, Molecular Devices). IC50 was calculated as using Prism 9 (GraphPad Software) nonlinear regression.

### Graphical illustrations

Graphical illustrations were made with Biorender.com.

## Figure Legends

**Fig S1.**
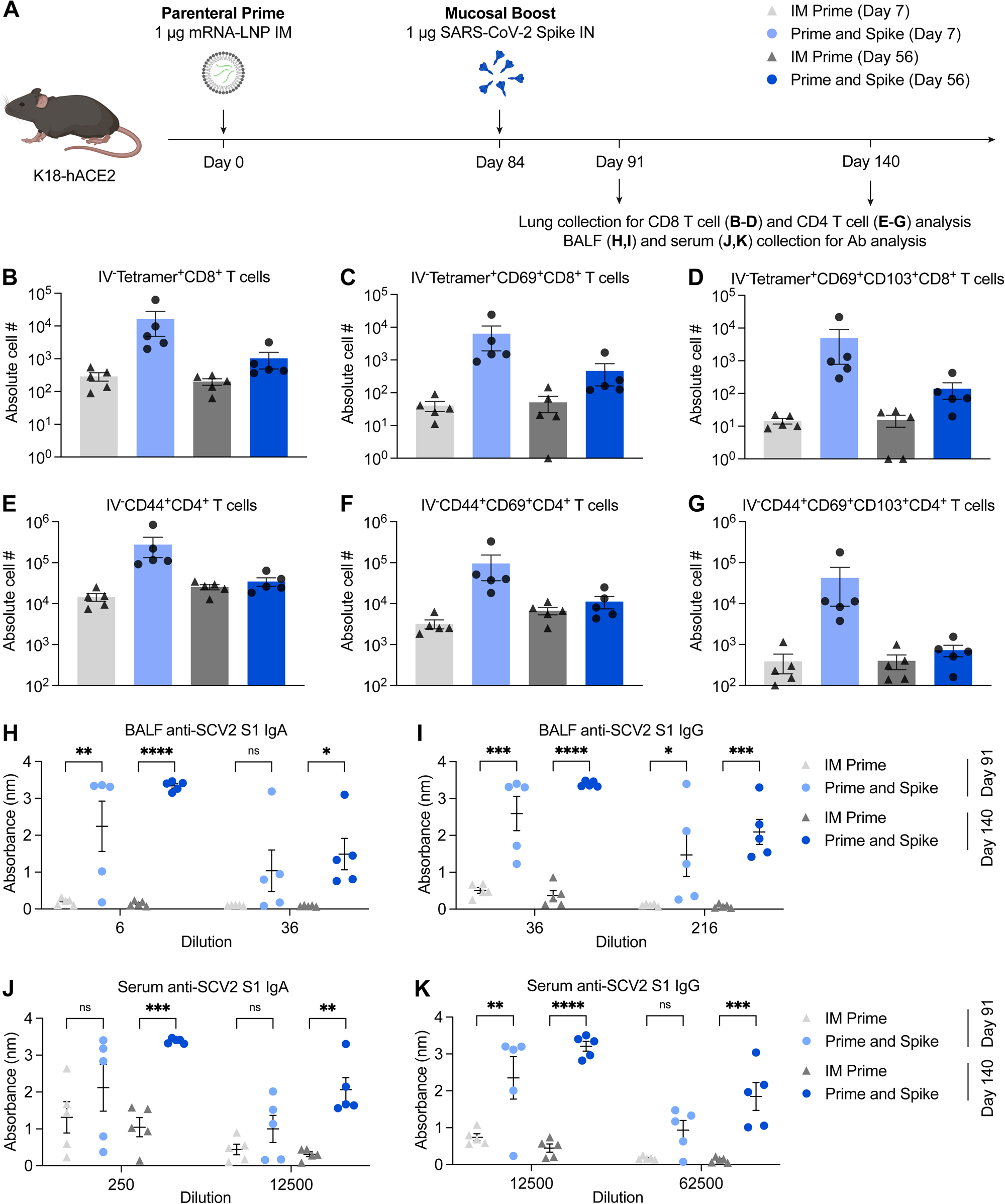
Delayed boosting with IN spike induces durable mucosal immunity. (**A**) Experimental schema: K18-hACE2 mice were IM primed with 1 μg of mRNA-LNP and IN boosted with 1 μg SCV2 spike 12 weeks post IM Prime. 7 and 56 days post boost, lung tissues were collected for T cell analysis by flow cytometry, and BALF and blood were collected for antibody measurement. (**B-D**) Quantification of total Tetramer^+^ CD8 T cells, CD69^+^CD103^-^ Tetramer^+^ CD8 T cells, or CD69^+^CD103^+^Tetramer^+^ CD8 T cells in lung tissues from IM Prime or Prime and Spike mice 7 and 56 days post boost. (**E-G**) Quantification of total activated, polyclonal CD4 T cells, CD69^+^CD103^-^ CD4 T cells, or CD69^+^CD103^+^ CD4 T cells in lung tissues from IM Prime or Prime and Spike mice 7 and 56 days post boost. (**H-K**) Measurement of SCV2 spike S1 subunit-specific BALF IgA (**H**), BALF IgG (**I**), serum IgA (**J**), and serum IgG (**K**) in IM Prime or Prime and Spike mice 7 and 56 days post boost. Mean ± s.e.m.; Statistical significance was calculated two-way ANOVA followed by Tukey correction (**H-K**); *P ≤ 0.05, **P ≤ 0.01, ***P ≤ 0.001, ****P ≤ 0.0001.

**Fig S2.**
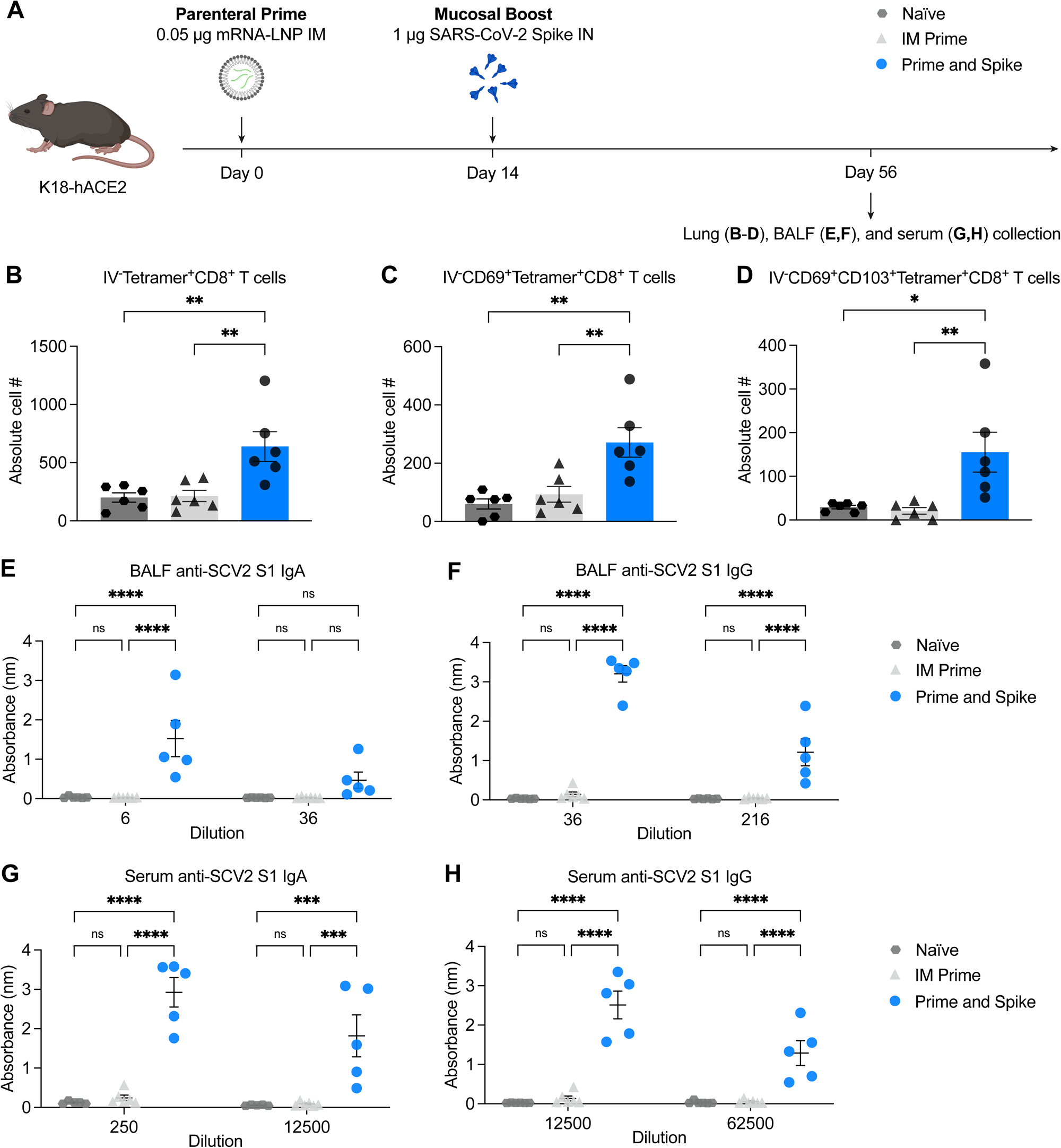
IN spike boost-induced protection against COVID-19 correlates with enhanced mucosal immunity. (**A**) Experimental schema: K18-hACE2 mice were IM primed with 0.05 μg of mRNA-LNP and IN boosted with 1 μg of spike IN 14 days post IM Prime. 6 weeks post boost, lung tissues were collected for CD8 T cell analysis by flow cytometry, and BALF and blood were collected for antibody measurement. (**B-D**) Quantification of total Tetramer^+^ CD8 T cells, CD69^+^CD103^-^Tetramer^+^ CD8 T cells, or CD69^+^CD103^+^Tetramer^+^ CD8 T cells in lung tissues from naïve, IM Prime, or Prime and Spike mice. (**E-H**) Measurement of SCV2 spike S1 subunit-specific BALF IgA (**E**), BALF IgG (**F**), serum IgA (**G**), and serum IgG (**H**) in naïve, IM Prime, or Prime and Spike mice. Mean ± s.e.m.; Statistical significance was calculated one-way ANOVA followed by Tukey correction (**B-D**) or two-way ANOVA followed by Tukey correction (**E-H**); *P ≤ 0.05, **P ≤ 0.01, ***P ≤ 0.001, ****P ≤ 0.0001.

**Fig S3.**
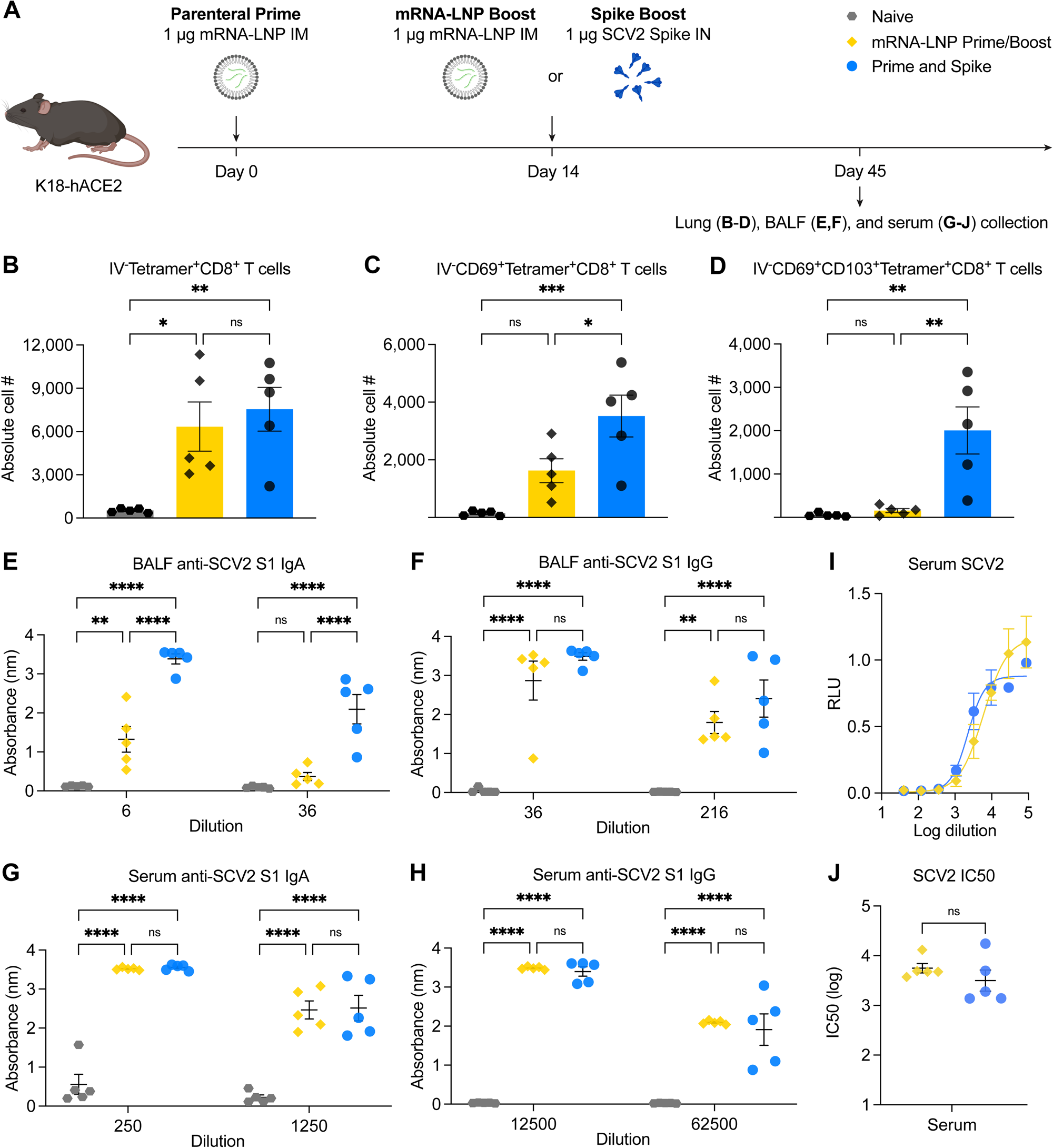
IN spike boosting elicits enhanced mucosal immunity with similar systemic humoral responses to IM mRNA-LNP boosting. (**A**) Experimental schema: K18-hACE2 mice were IM primed with 1 μg of mRNA-LNP, followed by boosting with 1 μg of mRNA-LNP IM, or 1 μg of SCV2 spike IN (IN Spike) 14 days post IM Prime. 45 days post boost, lung tissues were collected for T cell analysis by flow cytometry, and BALF and blood were collected for antibody measurement. (**B-D**) Quantification of total Tetramer^+^ CD8 T cells, CD69^+^CD103^-^Tetramer^+^ CD8 T cells, or CD69^+^CD103^+^Tetramer^+^ CD8 T cells in lung tissues from naïve, mRNA-LNP Prime/Boost, or Prime and Spike mice. (**E-H**) Measurement of SARS-CoV-2 spike S1 subunit-specific BALF IgA (**E**), BALF IgG (**F**), serum IgA (**G**), and serum IgG (**H**) in naïve, mRNA-LNP Prime/Boost, or Prime and Spike mice. (**I**,**J**) Measurement of neutralization titer against SCV2 spike-pseudotyped VSV. Mean ± s.e.m.; Statistical significance was calculated one-way ANOVA followed by Tukey correction (**B-D**), two-way ANOVA followed by Tukey correction (**E-H**), or student’s t-test (**J**); *P ≤ 0.05, **P ≤ 0.01, ***P ≤ 0.001, ****P ≤ 0.0001.

**Fig S4.**
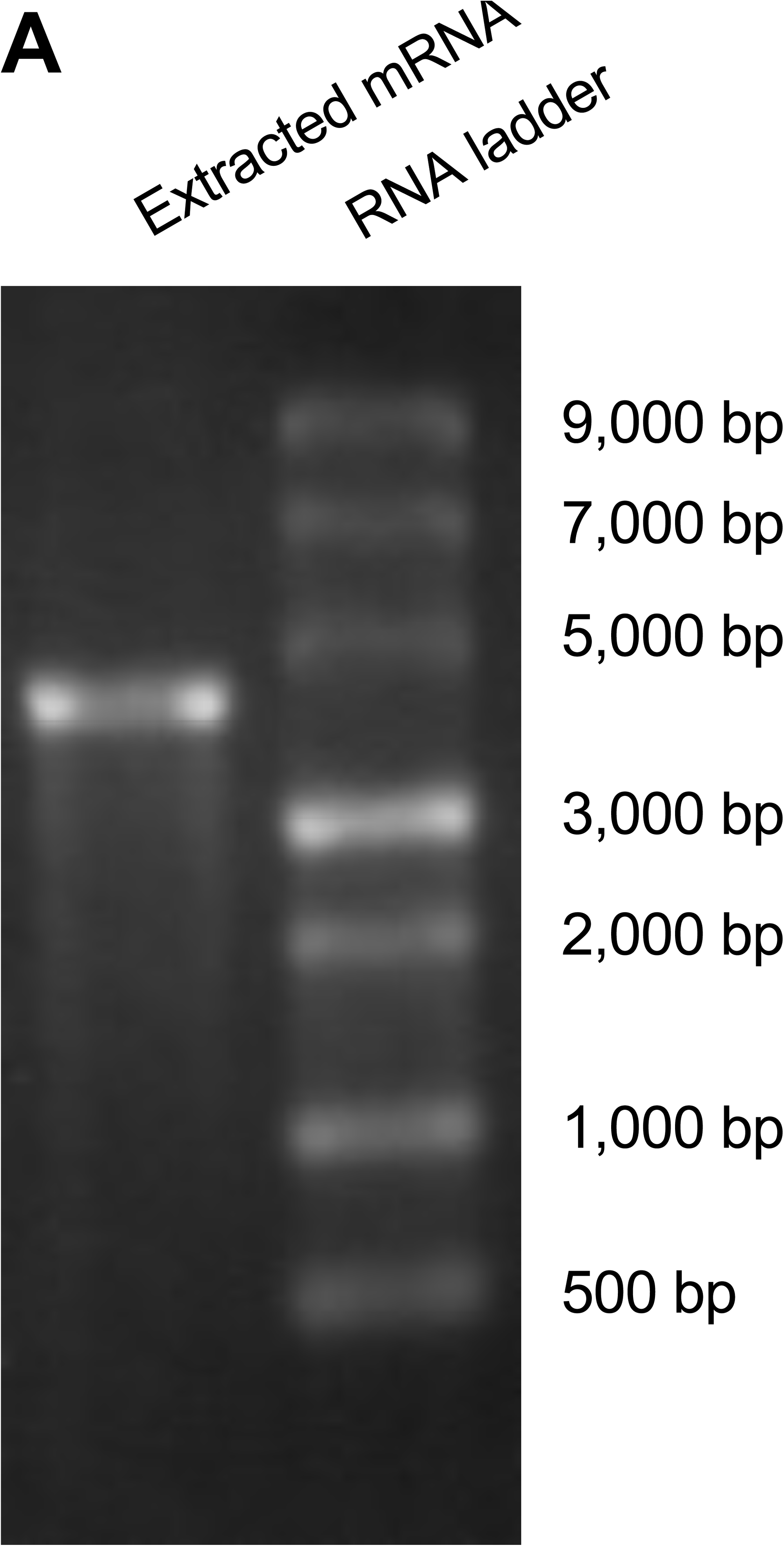
Confirmation of integrity of mRNA extracted from Comirnaty mRNA-LNP. (**A**) The length and integrity of extracted mRNA was analyzed using agarose gel electrophoresis. Extracted mRNA was mixed with SYBR Safe stain before being loaded onto a 1% agarose gel, let run in the TAE buffer, and imaged with a gel imaging system.

**Fig S5.**
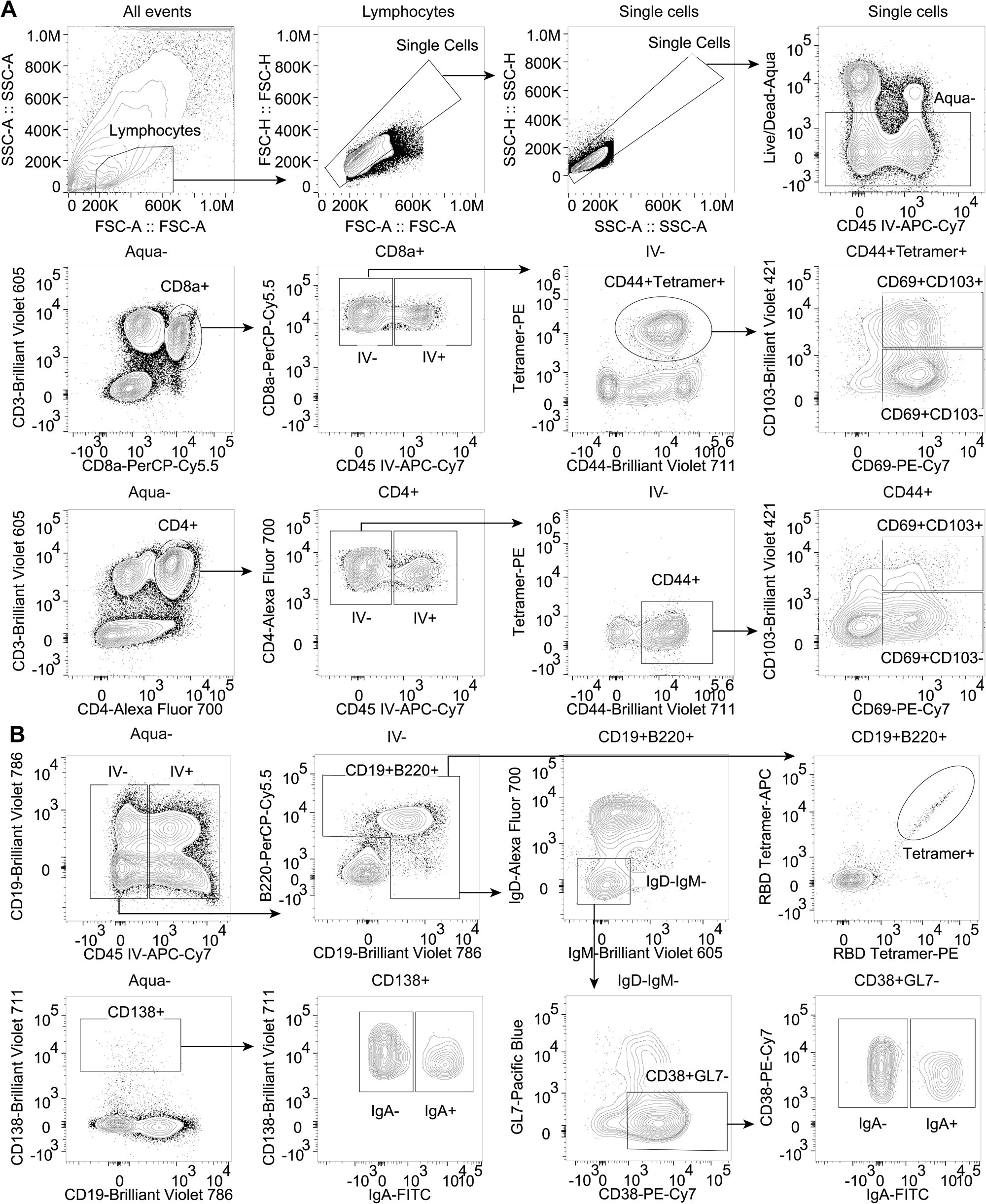
Gating strategies for analysis of extravascular adaptive immune responses in the respiratory tract. (**A**) Gating strategies to identify extravascular antigen-specific CD8 T cells and polyclonal activated CD4 T cells. (**B**) Gating strategies to identify extravascular antigen-specific and polyclonal B cell subsets.

## Acknowledgement

We thank M. Linehan for technical and logistical assistance. We thank Patrick Roberts and Bryan Cretella from the Yale Health Pharmacy for providing residual vaccine used in this study. We thank Barney Graham (NIH-VRC) for kindly providing VeroE6 cells overexpressing ACE2 and TMPRSS2. We thank the NIH Tetramer Core Facility for providing PE labelled SARS-CoV-2 S 539-546 tetramer (H-2K(b)). We thank Holly Steach for providing the B cell tetramer production protocol. We thank Craig Wilen and Jin Wei for technical expertise. We thank H.W. Suh for providing PACE polymers and A.S. Piotrowski-Daspit for helpful discussions. We also give special recognition of the services of Ben Fontes and the Yale EH&S Department for their on-going assistance in safely conducting biosafety level 3 research. This work was in part supported by the Fast Grant from Emergent Ventures at the Mercatus Center and 1R01AI157488. A.I. is an Investigator of the Howard Hughes Medical Institute. B.I. is supported by NIAID T32AI007517 and K08AI163493. T.M. is supported by NIAID T32AI007019. W.M.S, A.S., and M.R. are supported by UG3 HL147352 from NIH.

## Author contributions

B.I., TM, and A.I. conceived of and planned the project. B.I., T.M., A.S., L.Q., M.R., and M.P.H. performed experiments. B.I. and T.M. analyzed and interpreted data. R.J.H provided blinded histopathological analysis. H.D. carried out animal husbandry. W.M.S provided expertise and critical reagents. B.I., T.M., and A.I. wrote the manuscript and all authors reviewed and provided feedback on the manuscript.

## Corresponding authors

Correspondence and requests for materials should be addressed to B.I or A.I.

